# *Candida albicans* dispersed cells signify a developmental state distinct from biofilm and planktonic phases

**DOI:** 10.1101/258608

**Authors:** Priya Uppuluri, Maikel Acosta Zaldivar, Matthew Z Anderson, Matthew J. Dunn, Judith Berman, Jose Luis Lopez Ribot, Julia Koehler

**Affiliations:** The Division of Infectious Diseases, Los Angeles Biomedical Research Institute at Harbor-University of California Los Angeles (UCLA) Medical Center, CA; Division of Infectious Diseases, Children’s Hospital, Boston, MA; Department of Microbiology, The Ohio State University, Columbus, OH; Department of Molecular Microbiology & Biotechnology, Tel Aviv University, Ramat Aviv, Israel; Department of Biology and South Texas Center for Emerging Infectious Diseases, The University of Texas at San Antonio, San Antonio, TX

## Abstract

*Candida albicans* surface-attached biofilms are sites of amplification of an infection through continuous discharge of cells capable of initiating new infectious foci. Yeast cells released from biofilms on intravenous catheters have direct access to the bloodstream. We previously reported that dispersed cells are largely lateral yeast cells that originate from the hyphal layers of the biofilm. Compared to their planktonic counterparts, these biofilm-dispersed yeast cells displayed enhanced virulence-associated gene expression and drug resistance. Little is known about the molecular properties of dispersed cells. We found that the inducer of dispersal, *PES1*, genetically interacts with the repressor of filamentation, *NRG1*, in a manner that supports a genetic definition of dispersed cells as yeast. We combined a flow biofilm model with RNA sequencing technology, to identify transcriptomic characteristics of freshly dispersed yeast cells versus biofilms or age-matched planktonic yeast cells growing in glucose-rich medium. Dispersed cells largely inherited a biofilm-like mRNA profile but with one stark difference: dispersed cells were transcriptionally reprogrammed to metabolize alternative carbon sources, while their sessile parents expressed glycolytic genes, despite exposure to the same nutritional signals. Our studies hence define dispersal cell production as an intrinsic step of biofilm development which generates propagules capable of colonizing distant host sites. This developmental step anticipates the need for virulence-associated gene expression before experiencing the associated external signals.

## Introduction

From the perspective of a pathogen, detachment from an established niche for entry into the blood stream enables progression in the host. This niche could be any site in the body where the organism silently colonizes as a commensal; or it could be the layers of a drug resistant biofilm growing on biotic surfaces (e.g. oral or urogenital mucosa) or abiotic platforms (indwelling catheters) in hospitalized patients.

The human commensal *C. albicans* is also the most frequently isolated human fungal opportunistic pathogen. Disseminated candidiasis carries unacceptably high crude and attributable mortality rates – about 40-60%, despite antifungal drug treatment ^1,2^. *C. albicans* is unique for the ease with which it switches between growth forms in the host, between budding yeast and filamentous pseudohyphae and hyphae. The switch from yeast to the hyphal form increases its ability to adhere, invade and sustain a community of sessile cells. Decades of research have elucidated the regulation of hyphal morphogenesis and its relationship with pathogenesis ^3–5^. For example, candidalysin, a toxin of human cells produced by *C. albicans*, is formed only during hyphal growth^6^.

Biofilms, an important mode of growth for *C. albicans* as for many other microorganisms, are predominantly composed of hyphal cells. Hyphal filaments constitutively produce lateral yeast cells on their subapical segments, while apical segments continue to extend as filamentous cells. Lateral yeast cells are formed *in vivo*, because filamentous cells are always accompanied by yeast cells in organs of hosts with invasive candidiasis^7,8^. Similarly, biofilm-associated hyphae continuously release lateral yeast cells throughout the biofilm growth cycle^9^. In a clinical setting, such dispersed cells released from biofilms growing on indwelling catheters or from infectious foci that access the blood stream, are the cells that disseminate throughout the host and initiate distant foci of infection.

Current understanding of lateral yeast production and biofilm dispersal is limited. Earlier we found that yeast arising from biofilms adhered better to mammalian cells, were more resistant to azole drugs and displayed higher virulence in a disseminated mouse model of candidiasis when compared to free-living planktonic *C. albicans* yeast cells, ^9^. Thus, dispersal of cells from biofilms appears to be a distinct developmental phase of the fungus, in which cells are primed for invasion of the host.

To date, only one molecular regulator, Pes1, that controls production of lateral yeast from hyphae and induces biofilm dispersal, is known^9–11^. Pes1 regulates lateral yeast cells without changing the hyphal morphology or affecting biofilm architecture. It is essential for sustained candidiasis^7^. Curtailing lateral yeast cell release from tissue-invading hyphae by depleting *PES1* significantly reduced dissemination and virulence in mice^8^. The goal of this work was to define the molecular characteristics of lateral yeast cells released from biofilms.

Here, we used RNA sequencing to delineate biofilm dispersed cell-specific molecular signatures, and to identify gene expression patterns that might set the dispersed cells apart from their parent biofilms or their planktonic counterparts. We found that the transcriptional landscape of dispersed yeast largely overlapped with that of their parent hyphal biofilms, rather than with the morphologically similar, planktonic yeast sister cells. However, despite arising from the same nutritional milieu, dispersed and biofilm cells also exhibited striking contrasts in their metabolic profiles. We found that the key regulator of the transition from hyphal to lateral yeast formation, *PES1* is required for the yeast form, and reveals that *PES1* is the primary link between biofilm-contaminated catheters and disseminated candidiasis, using a jugular vein catheterized mouse model. In summary, we define molecular characteristics of yeast dispersed from biofilms and show that they are in a developmental phase distinct from the biofilm or planktonic phase, and are specifically equipped for immediate infection in the host.

## Results

### Transcript profiles of biofilm dispersal cell populations strongly correlate with those of sessile cells

To understand the molecular characteristics of *C. albicans* cells freshly released from an established biofilm, we collected yeast cells spontaneously released from biofilms, using the simple flow biofilm model ^12,13^. Age-matched planktonic and biofilm cells were recovered at 24 hours, and transcript profiles of the three populations were delineated through RNA-Seq.

We first compared differences between biofilm and planktonic cells. A total of 1524 genes exhibited significantly altered expression, using a false discovery rate (adjusted p-value) of <0.05, between the two cell types. The molecular profile of sessile versus free-living cells was reported previously in several genome-wide studies, as reviewed in ^14^. Our analysis of biofilm cells largely replicated and thereby underscored these findings^15,16^, concurrently validating the procedure (Fig. 1 and S1).

**Figure 1.**
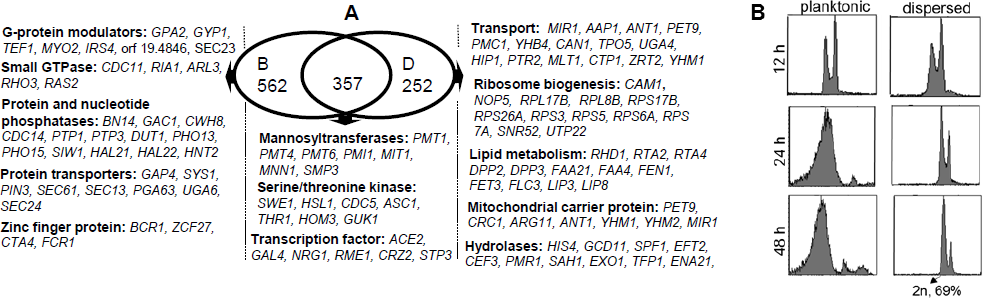
Venn diagram of significantly upregulated genes in biofilm and biofilm dispersed cells at 24 h, compared to planktonic cells. Gene expression profiles of *C. albicans* biofilm and biofilm dispersed cells grown in YNB with 2% glucose at 37 oC, were each compared individually to age matched planktonic cells. The Venn diagram displays total number of genes upregulated in each growth condition, and those common between biofilms and dispersed cells. The diagram also shows some genes and their functional categories (additional to those mentioned in the body of the manuscript) that are elevated in each group.

Next, we compared the biofilm-dispersed cells to their age-matched planktonic sisters. A total of 963 genes showed statistically significant differences between the two conditions (p<0.05). Out of the 609 genes upregulated > 1.6 fold in dispersed cells, 357 (∼59%) were also upregulated in a mature biofilm compared to planktonic cells (S1). Regulatory patterns of dispersed cells substantially reflected those of their parent biofilms with respect to genes involved in methionine biosynthesis (*MET1*, *MET2*, *MET3*, *MET10*, *MET15*, *MET16*, *SAM2*, *SAM4, SSU1*)^15^, aromatic amino acid metabolism (*ARO2*, *ARO3*, *ARO4*, *ARO8*, *ARO9*), ergosterol biopsynthesis (a majority of *ERG* genes), adhesion (*ALS3*, *ALS6, ECM33*)^17,18^, small molecule efflux and drug resistance (*MDR1, QDR1*)^19,20^, chitin synthesis (*CHS1*, *CHS2*, *CHS8*, *CHT1*, *CHT2*)^21^, glycerol biosynthesis (*GPD1*, *GPD2*, *RHR2*)^22^, almost all ribosome biogenesis genes^15,23^, and many others defined previously as biofilm regulated^14^. Expression of the yeast-specific gene *YWP1*^24^ was two-fold higher in dispersed cells than in biofilms, and higher in both compared to planktonic cells.

The patterns of expression of virulence genes such as Secreted Aspartyl Proteases (*SAP*) and lipase genes in dispersed cells appeared similar to those of the biofilms. Compared to planktonic cells, a majority of the *SAP* family genes, *SAP-3*, *4, 6, 8*, and *9* were upregulated in dispersed and biofilm cells, while *SAP4* and *SAP10* were also elevated in biofilm cells. Similarly, *LIP3* and *LIP8* were upregulated in dispersed cells, with *LIP4* additionally elevated in biofilm cells. Almost none of the *SAP* or lipase family genes were expressed in planktonic cells, similar to previous reports^25^.

### Dispersed yeast cells express genes associated with the hyphal form, like parent biofilms

In this study (and as reported by us previously)^9^, spontaneous dispersal cells from undisturbed biofilms were predominantly in an unbudded, elongated yeast form. We next asked if these cells are released from their hyphal mothers at a specific cell cycle stage. DNA content of the dispersed yeast was analyzed using flow cytometry, and compared to planktonic yeast cells at different time points of growth. As seen in Fig 2, at 12 h the DNA signal of both planktonic as well as dispersed cells was 4n, suggesting enrichment for G2/M cells, with a substantial fraction between 2n and 4n, indicating many cells in S-phase with ongoing DNA replication. At later time points (24 and 48 h), the DNA content of the planktonic cells shifted to the 2n state, with a lower percentage in S phase. At 48 h, almost all the planktonic cells and a majority (69%) of the dispersed cells were found in the 2n (unbudded) state, most likely reflecting stationary phase. ^26^

**Figure 2.**
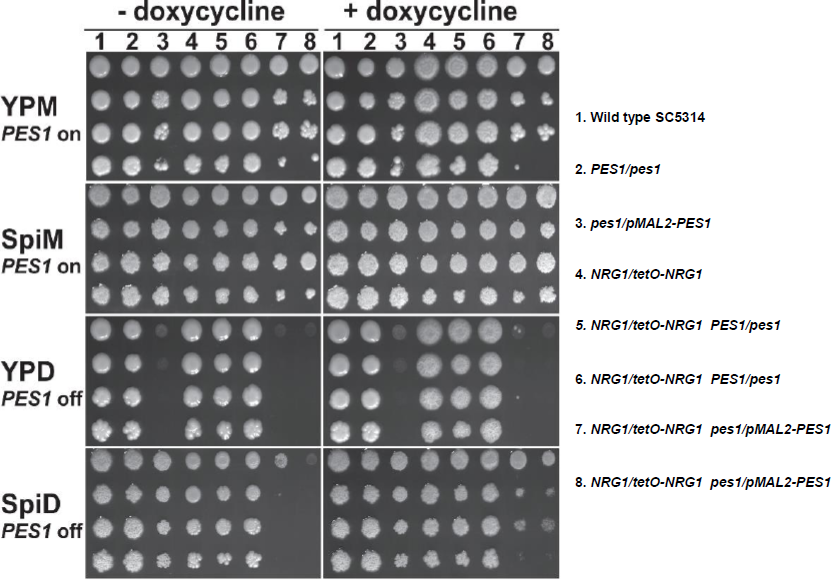
Analysis of cellular DNA by fluorescence flow cytometry. *C. albicans* planktonic and flow-biofilm dispersed cells at various time points of growth (12 h, 24 h and 48 h) were collected, and their DNA was labeled with propidium iodide. The cells were sorted in a flow cytometer to determine the stage (2n or 4n) of DNA replication. Over 69% of both planktonic and dispersed cells were found to in the unbudded, 2n stage of growth.

Despite their yeast morphology, and the expression of two predominantly yeast-specific genes (*YWP1* and *RHD3*)^27^, several genes upregulated in the dispersed population are typically expressed in hyphal, rather than yeast, cells. These include *DDR48, PHR2*, *ASC1, SUN41, ACE2, CDC5, CHA1, PCL5, PGA13, PGA26, PMR1, PMT1, PMT4, PMT6, RBR1, UTR2, YNK1*. To further characterize the dispersed cells, we developed two separate biofilm assays using two independent *C. albicans* strains in which the hyphal wall protein *HWP1* and yeast wall protein *YWP1* genes were fused with red fluorescent protein, *RFP.* These reporter strains showed that ∼33% of dispersed yeast expressed hyphal-specific *HWP1-RFP.* The *YWP1-RFP* control was expressed in ∼64% of dispersed yeast. These findings are striking for the substantial dissociation between the growth form of dispersal cells and their expression of cell-type specific cell wall genes.

Important genes involved in cytokinesis of hyphal cells, like those coding for actomyosin ring assembly protein Iqg1, were increased in both biofilm and dispersed cells. Also elevated were genes needed for ingression of the plasma membrane into the bud neck and primary septum formation such as *CHS2* (in dispersed cells), and *CYK3* and *HOF1* (in biofilm cells)^28,29^. Finally, genes encoding the mitotic exit phosphatase Cdc14 that modulates Chs2 trafficking and Iqg1 stability^28^ and a kinase that regulates cytokinesis in *C. albicans* filaments- *CDC5*, cell separation via the daughter cell specific transcription factor *ACE2*, were upregulated > 3 fold in both dispersed and biofilm cells, versus planktonic cells. Taken together, these results suggest that cytokinesis and cell separation are upregulated in biofilm hyphal cells, likely in preparation for release of daughter dispersed yeast cells.

### The dispersal cell inducer, *PES1*, is essential in cells overexpressing *NRG1*

Since the transcriptional profile of dispersed cells most closely resembled that of hyphal biofilm cells, while their morphological appearance corresponded to yeast, we examined dispersed cells by another criterion of cell type: essentiality of the inducer of lateral yeast growth, *PES1*. *PES1*, whose homologs are essential in all eukaryotes studied including the model yeast *S. cerevisiae*, is required for *C. albicans* yeast cell growth, but hyphae tolerate its depletion from repressible promoters^7^. Overexpression of *PES1* in biofilms induces increased lateral yeast production and dispersal from hyphal layers of the biofilm^9^. Overexpression of *NRG1*, a negative regulator of filamentation, also results in significantly increased production of dispersed cells^11^ from a predominantly hyphal biofilm. Here, we found that *PES1* message was elevated >1.7 fold in dispersed cells compared to biofilm cells, while *NRG1* expression trended lower (1.4 fold, p value 0.03, p-adj = 0.08) in dispersed versus the biofilm cells.

The genetic relationship between *NRG1* and *PES1* with respect to dispersal cell production was investigated in an *in vitro* static biofilm model. For this purpose, a *pes1/pMAL2-PES1 NRG1/tetO-NRG1* strain was constructed and compared with strains containing similar mutations in *PES1* or *NRG1* alone. In the presence of maltose, the *pes1/pMAL2-PES1* strain produced a biofilm with many lateral yeast cells (Fig. 4 A), while in glucose (during depletion of *PES1*) lateral yeast cells were significantly decreased (Fig. 4 B). The metabolic activity of the biofilms measured by XTT assay, was at least 33% higher in biofilms formed in maltose than in those formed in glucose, presumably reflecting a higher total number of cells when *PES1* expression permitted production of lateral yeast from biofilm hyphae. Nrg1, known to repress hyphal growth, repressed biofilm formation: *NRG1* upregulation abrogated biofilm formation (Fig. 4 C), while abundant hyphae and a robust biofilm were produced during *NRG1* repression (YNB+DOX, Fig. 4 D). *PES1* up- or downregulation when *NRG1* was overexpressed in the absence of doxycycline, did not reverse the biofilm defect; cells overexpressing *NRG1* remained in the yeast form regardless of the *PES1* expression level (Fig. 4 E, F).

**Figure 3.**
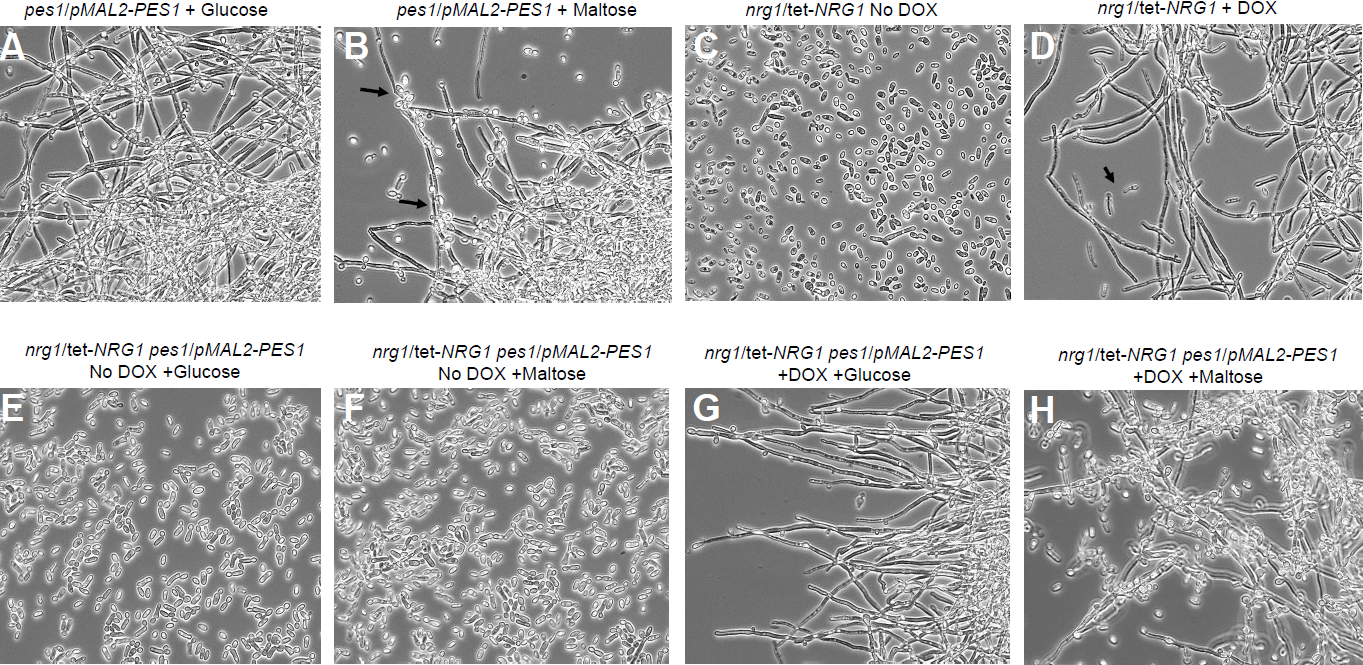
Epistasis study of *PES1* with *NRG1* in planktonic conditions. Serial dilutions of strains were spotted onto YP or Spider medium with Glucose or Maltose 2% as carbon source, without or with DOX 5 µg/ml. (1) Wild type SC5314; (2) *PES1/pes1*, JKC619; (3) *pes1/pMAL2-PES1*, JKC673; (4) *NRG1/tetO-NRG1*, SSY50-B; (5) *NRG1/tetO-NRG1 PES1/pes1*, JKC869; (6) *NRG1/tetO-NRG1 PES1/pes1*, JKC870; (7) *NRG1/tetO-NRG1 pes1/pMAL2-PES1*, JKC993; (8) *NRG1/tetO-NRG1 pes1/pMAL2-PES1*, JKC996. Plates were photographed after 3 days of incubation at 37°C (Spider) or 30°C (YP).

**Figure 4.**
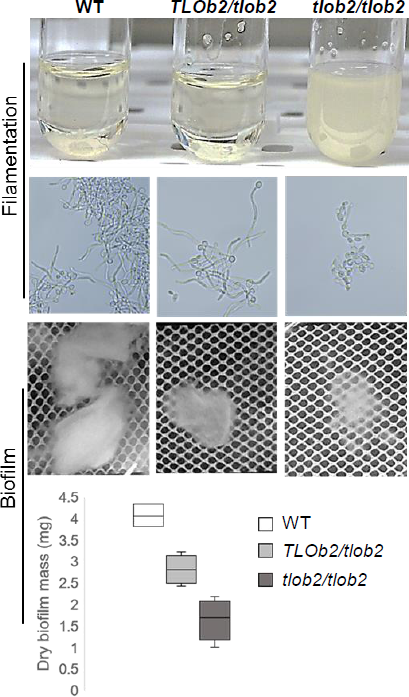
Epistasis study of *PES1* with *NRG1* under biofilm conditions. Biofilms were developed overnight in YNB broth in the presence of glucose or maltose as the carbon source, and in the presence or absence of DOX. The following strains were used: *pes1*/*pMAL2-PES1, nrg1*/tet-*NRG1*, and *nrg1*/tet-*NRG1 pes1*/*pMAL2-PES1*. Biofilms were visualized by light microscopy. Arrows in panel B and D point to the lateral yeast cells.

In contrast, lateral yeast production was significantly impacted by *PES1* induction or repression in *NRG1-*repressed hyphal biofilms. Downregulation of *PES1* abrogated lateral yeast formation and biofilm dispersal (Fig. 4 G), while induction of *PES1* expression in *NRG1*-downregulated biofilm cells resulted in prolific lateral yeast production, a 3-fold increase in biofilm dispersal (data not shown), and a 36% increase in biofilm metabolic activity as monitored by XTT assay compared to *PES1/NRG1* repressed biofilms (Fig 4 H). When *pMAL2*-*PES1* expression was induced during *NRG1* downregulation, lateral yeast cells significantly increased. These findings supported the definition of dispersal cells produced on biofilms as yeast. They also showed that for dispersal cell production, *PES1* is epistatic to *NRG1*.

In control experiments, we tested genetic relationships of *NRG1* and *PES1* for growth under yeast- or hyphae-inducing conditions. Consistent with previous findings, a glucose repressible *pes1/pMAL2-PES1* strain failed to grow at 30° C in YPD medium (yeast extract medium with 2% glucose), yet grew at 37° C in Spider medium containing glucose as the carbon source (Fig 3, Lane3). *NRG1/tetO-NRG1* strains with wildtype *PES1* grew in all media, with colonies comprised primarily of yeast cells during *NRG1* upregulation (- DOX), and primarily of hyphal cells during *NRG1* repression (+ DOX, as previously established by others^30^, Lanes 4,5,6). When *PES1* was depleted in *NRG1*-overexpressing cells, growth ceased (-DOX, lanes 7, 8), indicating that these cells, like yeast cells that are wild type for *NRG1*, are *PES1* dependent. In contrast, when *PES1* was depleted in *NRG1*-repressed cells, these hyphal cells grew robustly (+DOX, lanes 7,8). Hence, during *PES1* depletion, growth depended on lifting genetically determined hyphal repression, indirectly supporting the idea that dispersed cells exhibit essential characteristics of yeast cells.

### Genes differentially regulated in dispersed cells have roles in zinc, iron and amino acid transport and pathogenesis

A total of 335 genes (167 up- and 168 down-regulated) were expressed differently in dispersed cells, compared to both biofilm and planktonic conditions (Listed in S2 sheet 2). The predominant group of genes upregulated only in dispersed cells represented functions in transport. Among high affinity transporters were those for zinc (*ZRT1, ZRT2*)^31^, lactate (*JEN1*, *JEN2*)^32^, iron (*FET3, CFL2, CFL4, CFL5*), amino acid- (*GAP1, GAP2, AAP1, UGA4, CAN1, ANT1, PTR2, OPT1, OPT8*), and mitochondria-associated transport (*YHM1*, *YHM2*, *PET9, NDE1, MIR1*, *CRC1*). In addition, genes associated with morphogenesis and biofilm growth had a distinct expression pattern in dispersed cells relative to biofilm and planktonic cells (e.g., *CSA1, YNK1, PGA26, GNA1, RBT1, ALS5*, *IFF4* and *UTR2)*. Other categories of upregulated genes included hydrolases (e.g., aspartic proteases, lipases etc.), and genes involved in lipid metabolism (Fig 1).

Of the 168 genes downregulated exclusively in the dispersed cells, 60 had unknown biological functions. While the remaining 108 genes were not readily categorized into functional groups, a number had functions either in mRNA binding, or coded for proteins with predicted activity in mRNA splicing via the spliceosome, E.g. *ZSF1*, orf19.285, *EXM2*, orf19.2261, orf19.7139, *FGR16*. Other downregulated dispersal cell-specific genes were predicted to function in retrograde transport from the endosome/ER to the Golgi, including *VPS41, VPS17, UFE1, MSO1, HNM3*, orf19.2333, orf19.5114. The significance of downregulation of these genes in dispersal cells remains to be determined.

### Telomere-associated (*TLO*) gene family expression shows stark differences between dispersed and sessile cells

The TLO gene family exhibited significant expression differences between biofilm cells, planktonic cells, and dispersed lateral yeast. This gene family, whose products are thought to be components of the transcription-regulating Mediator complex [ref], is comprised of 13 telomere adjacent genes (*TLO1-5*, *TLO7-13*, *TLO16*) ^33,34^ and one internal copy of the gene (*TLO34*). In dispersed cells, most *TLO* genes were downregulated at least four-fold relative to biofilm cells, and either equal in expression (*TLO4, TLO5, TLO7, TLO13*) or two-fold downregulated relative to planktonic cells (*TLO1, TLO2, TLO9, TLO10, TLO34*), respectively. In contrast, several *TLO* genes (*TLO1, TLO2, TLO5, TLO7, TLO8, TLO11, TLO13 and TLO16*) were most highly expressed in biofilm cells, with the most extreme case being *TLO2*, which was 7-fold higher in biofilms and 2 -fold higher in planktonic cells than in dispersed cells.

*TLO* genes are organized into three clades with multiple family members in the alpha and gamma clades and only one member, *TLO2* in the beta clade ^33^. We investigated the morphogenesis-, biofilm- and dispersed phenotypes of a mutant lacking *TLO2*, since expression of this gene most distinguished biofilm cells from dispersed and planktonic cells. The *tlo2-/-* strains were defective in filamentation (Fig 5): the cells remained uniformly suspended in liquid Spider medium, in contrast to the clumps of adherent filaments produced by the wild-type and heterozygous parent strains, which settled to the bottom of the tube. The *tlo2-/-* strain produced almost exclusively yeast cells (Fig. 5). Under conditions of static biofilm induction, *tlo2-/-* showed a significant decrease (∼60%) in biofilm growth, compared to the wildtype strain (Fig. 5) with the *TLO2/tlo2-* strain having an intermediate phenotype (∼31% decrease). Biofilms formed by the null mutant were fragile, and tended to disintegrate easily, as a consequence of the deficiency in developing scaffolding filaments. Given their defects in biofilm formation, we were not able to assess the dispersal cell phenotypes of *tlo2-/-* mutants. Further experimentation, e.g. with repressible *TLO2* alleles whose expression can be downregulated after a biofilm has formed, will be required to define the distinct roles of Tlo2 in biofilm-versus dispersal cells. Such experiments could answer the question which transcriptional targets of Tlo2 are differentially regulated in hyphal biofilm cells and their dispersal cell daughters.

**Figure 5.**
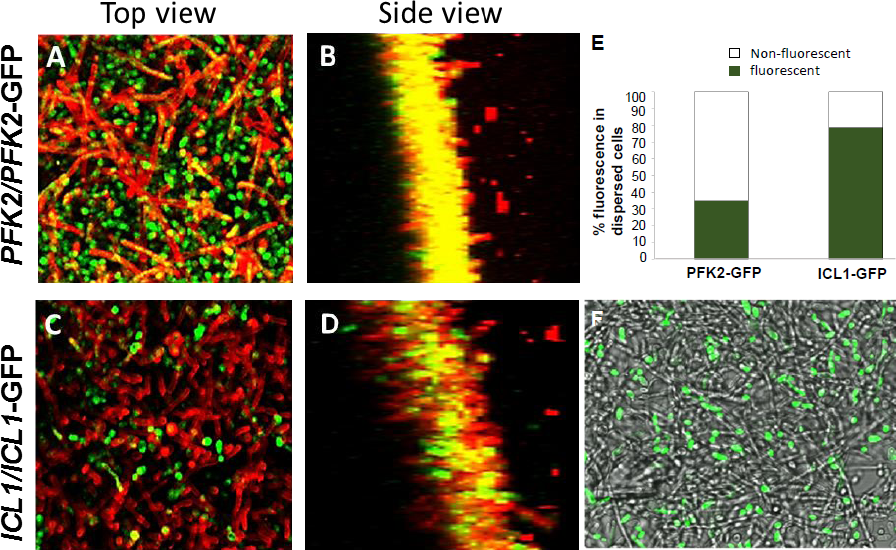
Filamentation and biofilm-forming capabilities of a *TLOb2* mutant of *C. albicans*. *Candida* strains (wildtype, *TLOb2* heterozygote and *tlob2/tlob2* homozygote) grown overnight in YPD at 30°C were inoculated into Spider media, and incubated at 37°C for 0, 2, 4 and 6 h. Aliquots from each strain were assessed for cell phenotype by microscopy. Biofilms for these strains were also developed under static conditions for 48 h, visualized microscopically, and their dry weight measured for biomass. The *tlob2-/-* homozygote showed a drastic reduction in both filamentation and biofilm formation compared to the wildtype strain. The heterozygote displayed normal hyphal development, but the hyphae did not clump together as was seen in the wildtype samples. The *tlob2/TLOb2* heterozygote also had an intermediate defect in biofilm formation.

### Dispersed cells resembled planktonic cells with respect to carbon metabolism

The biofilm in this study was developed under continuous flow of fresh medium, rich in glucose (1%). Biofilm cells expressed much higher levels (3-9 fold) of several key genes encoding the glycolysis pathway enzymes (*GAL4*, *PGI1*, *PFK1, PFK2, FBA1, PGK1, ENO1, CDC19*) relative to 24 h planktonic cells. Dispersed cells released from these same biofilms displayed the opposite pattern of expression (S2): all glycolysis genes were repressed and core genes required for alternative carbon metabolism were highly upregulated. This pattern of gene expression in dispersal cells resembled that of planktonic cells, wherein elements of the gluconeogenesis pathway - *PCK1* and *FBP1* - had six and three-fold increases, respectively, relative to biofilm cells. Genes encoding enzymes of the tricarboxylic acid (TCA) pathway presented the greatest differences: all were upregulated > 10 fold in dispersed as well as planktonic cells, versus sessile cells (S2, sheet 1). Importantly, *ICL1*, which encodes a major enzyme of the glyoxylate pathway, was upregulated >20 fold in planktonic cells and 14-fold in dispersed cells relative to mature biofilms. Thus dispersed cells sharply differed from their hyphal mothers, while resembling planktonic cells, in their upregulation of alternative carbon metabolism-related genes.

### Expression patterns shifted in lateral yeast cells before their release from the biofilm

To visualize the disparity of carbon metabolism gene expression between biofilm and dispersal cells at the protein level, we used strains expressing *GFP*-tagged *PFK2* (a glycolysis gene) or *ICL1* (a glyoxylate cycle gene). Biofilms were then developed under the same growth conditions as those of cells analyzed by RNA seq. After 24 h, biofilms were stained using Concanavalin A (ConA, a lectin that binds to mannans of the fungal cell wall), and visualized by confocal microscopy. A top-down view of *PFK2*-GFP biofilms displayed a high intensity of green fluorescence (with yellow streaks in hyphae due to overlap with red ConA signal), suggesting that biofilm cells were performing glycolysis. Merging signals of the side view of the biofilm showed overlapping signals, indicating a potential for glycolytic activity in hyphae stained by ConA (Fig 6 A, B, also see S3 slide 1 for additional alternative images). In contrast, only a small proportion of cells in the *ICL1*-GFP biofilms showed green fluorescence (Fig 6 C, D), suggesting that the glyoxylate cycle was largely inactive in the biofilms.

**Figure 6.**
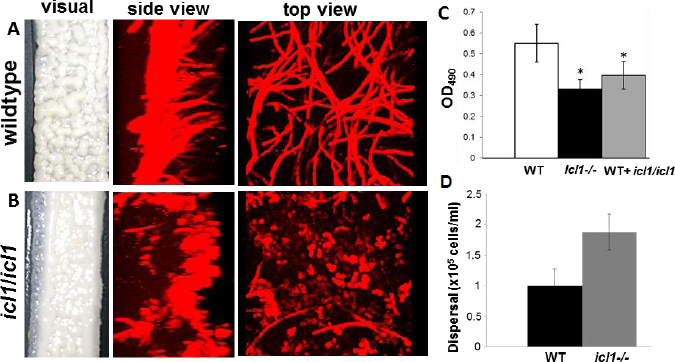
Determination of the metabolic preference of biofilm and biofilm dispersed cells. *C. albicans* cells harboring GFP tagged glyoxylate pathway gene *ICL1* (*ICL1/ICL1-GFP*) and GFP tagged glycolytic gene (*PFK2/PFK2-GFP*) were allowed to develop biofilms individually, for 48 h. The biofilm cell walls were stained with ConA (red), and z-stacks were collected under two wavelengths 488 (GFP), and 594 (ConA). Top and side scatter images reveal that top-most hyphal layers of the biofilm express Pfk2 (A, B), rather than Icl1 (C,D). In contrast, the cells dispersed from the biofilms expressed Icl1 at a significantly higher frequency than Pfk2 (E). This can be visualized by overlaying a bright-field and GFP fluorescence image of the biofilm, which show brightly fluorescent yeast cell on top of a mat of non-fluorescent *ICL1-GFP* biofilms (F).

Interestingly, dispersed cells expressed *ICL1* at a significantly higher rate than *PFK2* (Fig 6 E). The dispersed cells from the *PFK2-GFP* biofilms displayed an overall dim fluorescence, and only about 35% of cells showed a clear fluorescent signal. In contrast ∼ 78% of the dispersed cells from the ICL1*-GFP* biofilms fluoresced bright green. A merged bright-field - GFP fluorescence image of the *ICL1-GFP* biofilm surface, clearly demonstrated that GFP fluorescence was evident only in lateral yeast cells, poised for release from the hyphal surface, and not in the hyphal matt (Fig. 6 F).

Since *ICL1* was not expressed at high levels in biofilms, we tested the hypothesis that *ICL1* was dispensable for biofilm growth under flow conditions. We compared a *C. albicans ICL1* mutant strain (*icl1*-/-) to the wildtype SC5314 strain. *C. albicans icl1*-/- appeared to form a significantly less robust biofilm by macroscopic inspection; confocal microscopy revealed that *ici1-/-* cells had a defect in filamentation under biofilm conditions (Fig 7 A, B). Consistent with previous reports, this defect did not appear under planktonic conditions^35^. Metabolic activity of the biofilms, measured by XTT assay, found that *icl1* mutants biofilms exhibited only ∼40% of the wild type metabolic activity (Fig 7 C). Thus, biofilms utilize the glyoxylate cycle for optimal metabolism.

**Figure 7.**
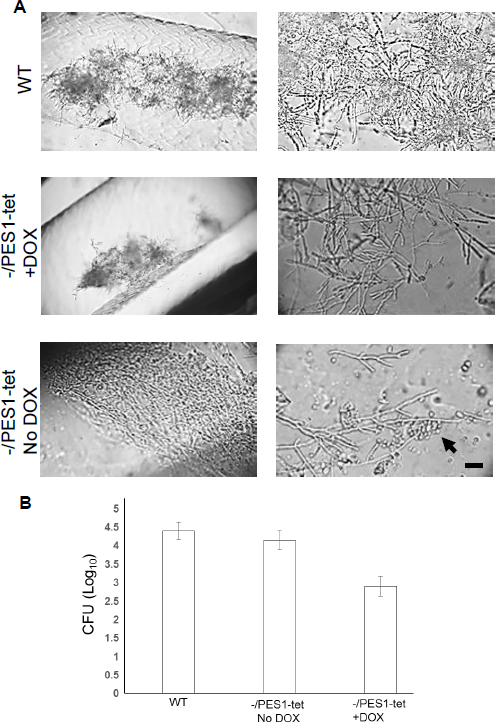
Impact of *ICL1* loss on biofilm formation. *C. albicans* wild-type and *icl1/icl1* strains were made to develop biofilms for 48 h. Macroscopic and microscopic observation (biofilms stained with ConA and imaged by CSLM) revealed that the mutant biofilms were less robust than the wildtype (A,B). This defect was confirmed by XTT assay that measures biofilm metabolism that can be read at OD490 (C). Nonetheless, the dispersal from *icl1* mutant biofilms was higher than the wildtype biofilms (D).

A mixed biofilm containing SC5314 and *icl1-/-* cells also demonstrated a significant reduction in biofilm growth (p<0.05).

Yeast cell dispersal from *icl1-/-* biofilms was two-fold increased over that of the wildtype biofilms (Fig 7 D). Thus, *ICL1* contributes to the metabolic activity of sessile growth; however, its detection was perhaps below the limit of detection under the experimental conditions used for RNAseq. Given this paradox, we dismantled the biofilm, inverted it and imaged its underside, as well as the thin basal layer of cells still attached to the silicone material. Here, we found that several patches of the inner-most layers of the biofilm showed an *ICL1-GFP* signal, indicating expression of Icl1 in this environment (S3 slide 2). Taken together these results suggest that a biofilm unit harbors subpopulations of cells, which have their own unique gene expression pattern. This was shown by *ICL1* expression within the deeper layers of the biofilm, where the cells experience substantial nutrient limitation. All the more striking was *ICL1* expression in lateral yeast cells, at the surface of the biofilm, where ambient nutrients are abundant. This implies that lateral yeast cells are likely to undergo metabolic reprogramming before release from the biofilm.

### Control of biofilm dispersal inhibits *C. albicans* dissemination in jugular vein-catheterized mice

We posit that cells released from biofilms growing on catheters are released into the blood stream, leading to candidemia and potentially to disseminated candidiasis. Thus, we asked if modulation of dispersal from biofilms *in vivo* could control biofilm-mediated invasive disease. Lateral yeast produced from hyphal layers of biofilms are the cell population that disperse. Since *PES1* is a major regulator of lateral yeast production and biofilm dispersal, we inoculated catheters intraluminally, with a *pes1/tetO-PES1* strain [ref Shen et al, 2008, paper about PES1]. In this strain, one allele of *PES1* is deleted while expression of the other is controlled by a 7xtetO regulatory sequence. In the absence of doxycycline (DOX), *PES1* is overexpressed constitutively[ref Shen et al, 2008, paper about PES1]; addition of doxycycline (DOX) in the growth medium suppresses *PES1* expression and drastically depletes the *PES1* message [ref Shen et al, 2008, paper about PES1]. This strain first was incubated for 90 minutes in the absence of DOX in YNB growth medium, and then introduced into catheters in YNB medium in the presence or absence of 25 uM DOX. As a control, wild-type *C. albicans* germ tubes were inoculated in YNB into independent catheters.

After 2 days, mice were sacrificed and the catheters were recovered aseptically and imaged. The extent of dissemination was measured by dissecting and homogenizing the kidneys, and plating for CFU on solid media. Microscopy of the biofilms found that all catheters harbored biofilms containing hyphal filaments; catheters in which *PES1* was overexpressed (in the absence of DOX) produced a high number of lateral yeast cells from the hyphal filaments (Fig 8). The extent of dissemination and kidney colonization was comparable between these catheters and wildtype biofilm-containing catheters. By contrast, biofilms in catheters treated with DOX to repress *PES1* lacked lateral yeast cells, and the kidneys of these mice showed a 1.5 log decrease in *C. albicans* dissemination and kidney colonization (Fig 8).

**Figure 8. Biofilm formation and dispersal in jugular vein catheterized mice.** Germinating *C. albicans* tetracycline regulatable *PES1* strain, *pes1/PES1-tet*, was instilled in the lumen of the catheters (full catheter volume) at a concentration of 5x10^6^ cells/ml. The cells existed in the catheters in YNB medium with and without doxycycline, for 3 days, after which the catheters were cut laterally and examined under a phase contrast microscope. While all the three strains developed biofilms in the catheters in the presence or absence of DOX, the Pes1-tet biofilm hyphae displayed increased lateral yeast production in the absence of DOX. Scale bar indicates 20 µm. Extent of biofilm-mediated dissemination in mice harboring the catheters containing the two strains, was also determined by measuring CFU counts in the kidney. In the presence of DOX, dissemination and kidney colonization was reduced 15 fold compared to the mice infected with catheters with wildtype or Pes1-tet without DOX.

## Discussion

Biofilm formation is the predominant mode of growth for most microorganisms in natural and clinical environments^36^. For many pathogenic organisms, biofilm dispersal plays a critical role in the transmission of cells between hosts, and in the propagation and spread of infection within a single host^37^. For example, *S. mutans* can detach from dental biofilms in a mother‘s mouth and be transmitted to an infant by direct or indirect contact^38^. *Candida albicans* cells are frequently detected along with heavy infection by *S. mutans* in plaque biofilms from early-childhood caries-affected children.^39^ Examples of intra-host spread encompass hospital-acquired pneumonia caused by bacteria detached from biofilms in a patient‘s endotracheal tube^40^; hematogenous spread of *C. albicans* into the eye, from an infected site post ureteric stenting and urinary sepsis^41^, or dissemination of *C. albicans* from biofilms on intravenous catheters as we modeled in this study. Hence, characteristics of dispersal cells are important to understand *C. albicans* pathogenesis.

We previously found that dispersed cells possess a number of phenotypic properties correlated with virulence^9^, compared to yeast cells growing as a free-living population, suggesting measurable distinctions between the molecular signatures of these two different populations of yeast cells. Thus, we initiated a global transcriptional profile investigation to identify the genes and pathways that might set the dispersed cells apart from their parent biofilm cells, or planktonic yeast cells which they resemble morphologically.

We utilized an *in vitro* model, where *C. albicans* biofilms were developed for 24 h under a continuous flow of fresh medium^12^. As reported previously, cells freshly dispersed from biofilms were in the unbudded form, similar to 24 h planktonically grown cells^9^. Furthermore, a majority of both populations of yeast cells possessed a single complement of chromosomal DNA after 24 hours (Fig.), indicating their basline characteristics may be sufficiently similar that comparisons of their gene expression profiles could be informative.

Differential expression of genes in our flow biofilms compared to age-matched planktonic cells, revealed similarities in more than 180 core genes known to have regulatory or non-regulatory roles in biofilm development^15,16,42,14^ (S1), validating this set of differentially regulated genes. Further, compared to planktonic cells, several genes important in adhesion were upregulated in dispersed cells to an extent analogous to the biofilm. This provided a molecular basis for our previous observation that dispersed cells are highly virulent^9^.

Similar to biofilm cells, and unlike planktonic cells, dispersed cells also exhibited elevated expression of the virulence-associated *SAP* genes. Expression of the *SAP* gene family remains at a low, steady state during planktonic growth starting from exponential to well into the stationary phase^26^. In contrast, biofilm cells secrete select *SAP*‘s to high levels as biofilm-specific proteases (candidapepsins)^25^, and several of these genes were in fact also identified as highly expressed in biofilm- dispersed cells (*SAP3, SAP4, SAP6, SAP8, SAP9*), in the present study.

Given the similarity of gene expression patterns of dispersal cells to their hyphal biofilm mothers, we wished to define the cell type of dispersal cells beyond their yeast-like morphological appearance. To do this, we examined the genetic relationship between *PES1* and *NRG1* in these cells. *PES1* is known to be essential in *S. cerevisiae* and *C. albicans* yeast cells, while hyphae tolerate its depletion^7^, so that cells requiring *PES1* expression can be defined as yeast cells. *NRG1* is a known repressor of hyphal growth, and overexpression of *NRG1* from *tetO* in the absence of doxycycline locks *C. albicans* in the yeast form, while repression of one allele from *tetO* in the presence of doxycycline induces hyphal growth (ref). Previous work showed that dispersal from biofilms is induced by *NRG1* or *PES1* overexpression, while *NRG1*-repressed biofilms are defective in producing dispersal cells (ref.). We now find that *PES1* overexpression from *tetO* induced prolific production of dispersal cells from *NRG1*-repressed biofilms (Fig.); in contrast, *PES1* repression in these *NRG1*-repressed hyphal biofilms blocked dispersal cell formation (Fig. x). These findings support the characterization of dispersal cells as yeast.

Increased expression of adhesins, hyphal genes and secretory aspartyl proteases in dispersed versus planktonic cells suggest that the biofilm-released cells are primed during their development to attach to and invade host tissues. Thus, *C. albicans* induce genes representing factors required during or after tissue invasion before the actual invasion process is initiated.

In variable environments, microbes may enhance their fitness by predicting and preparing for a coming change from cues arising in current conditions ^43^. As an example, the gut bacterium *Escherichia coli* elicits a transcription response for hypoxia when shifted from growth at 30°C to 37°C^44^, because increase in temperature “signals” its arrival in the gut where oxygen will soon become scarce. Similarly, *C. albicans* responds to higher pH by expressing genes involved in iron and zinc uptake via an alkaline-induced transcription factor Rim101^45^. Since the two metals are less soluble at high pH, the fungus predicts metal starvation based on its current growth environment. We posit that *C. albicans* dispersal cells express virulence factors in an anticipatory manner to optimize their transition from a sessile lifestyle to a disseminative and invasive form.

In a striking discrepancy of expression profiles between biofilm hyphal mothers and dispersal yeast daughters, flow biofilm cells expressed a metabolic profile reflective of glycolysis while yeast dispersed from these biofilms strongly expressed elements of the gluconeogenesis and the glyoxylate pathway. Dispersed cells may anticipate needing alternative carbon metabolism genes as a pre-adaptive response, to access host sites in tissues that have low glucose availability.

Unlike the yeast to hypha transition, that requires a specific external trigger (such as heat, serum, specific nutrients, alkaline pH, etc); the hypha to lateral yeast developmental process occurs constitutively. In nutritionally rich or poor media^7,46,47^, in mouse kidneys^8^, or biofilms on catheters^9^, *C. albicans* hyphae always produce lateral yeast cells. Hence this process appears to reflect an intrinsic developmental program. A precedent of regulated developmental steps that differentiate a fungal daughter cell from its mother is mating-type switching in *Saccharomyces cerevisiae*, in which *ASH1* mRNA is transported exclusively into the daughter, and represses switching in the daughter but not the mother^48^. *ASH1* is also found in *C. albicans* daughter but not mother cells^49^. *C. albicans* has a She3-mediated system that transports selected transcripts into daughter cells of budding yeast (and hyphal tips), and one third of these transcripts have roles in hyphal development^50^. *TLO* genes which are broad transcriptional regulators are differentially upregulated in biofilm cells. Whether the mechanism underlying differential gene expression in a biofilm mother cell and its dispersal daughter consists of selective mRNA transport between the cytoplasm of these cells, or of reprogramming of gene expression after mitosis of the dispersal cell, remains to be determined in future work.

### Role of dispersed cells in pathogenesis

Our present data reveals the importance of *PES1* for *C. albicans* yeast growth, which has a significant direct impact on lateral yeast production and biofilm dispersal. Regulation of *PES1* could have significant consequences in clinical settings, where catheters often harbor biofilms with direct access to the blood stream. This was seen in our studies with jugular vein catheterized mice, where overexpression of *PES1* in biofilms growing in the lumen of the catheters resulted in enhanced dispersal and disseminated infection in mice, while repression of *PES1* resulted in abrogation of biofilm dispersal and a 15 fold decrease in *Candida* infection of distal organs.

Effective management of disseminated candidiasis lies in the discovery of novel inhibitors that can interfere with the broad intracellular and regulatory signals controlling cellular propagation from colonized niches such as biofilms. An assimilation of awareness gained from *in vitro* studies on biofilms, together with a better understanding of the molecular signatures of biofilm dispersal, will pave the way for control of biofilm-mediated disseminated diseases in humans.

## Material and Methods

### Strains and culture conditions

Stock cultures of all strains were stored in 15% glycerol at −80°C. Strains were routinely grown under yeast conditions (media at 30°C), in YPD (1% yeast extract, 2% bacto peptone, 2% glucose), or in filament inducing conditions - RPMI (Sigma, St. Louis, MO) with MOPS buffer or Spider medium. Strains used in this study, their origin and their construction are briefly listed in Table 1.

**Table 1.**
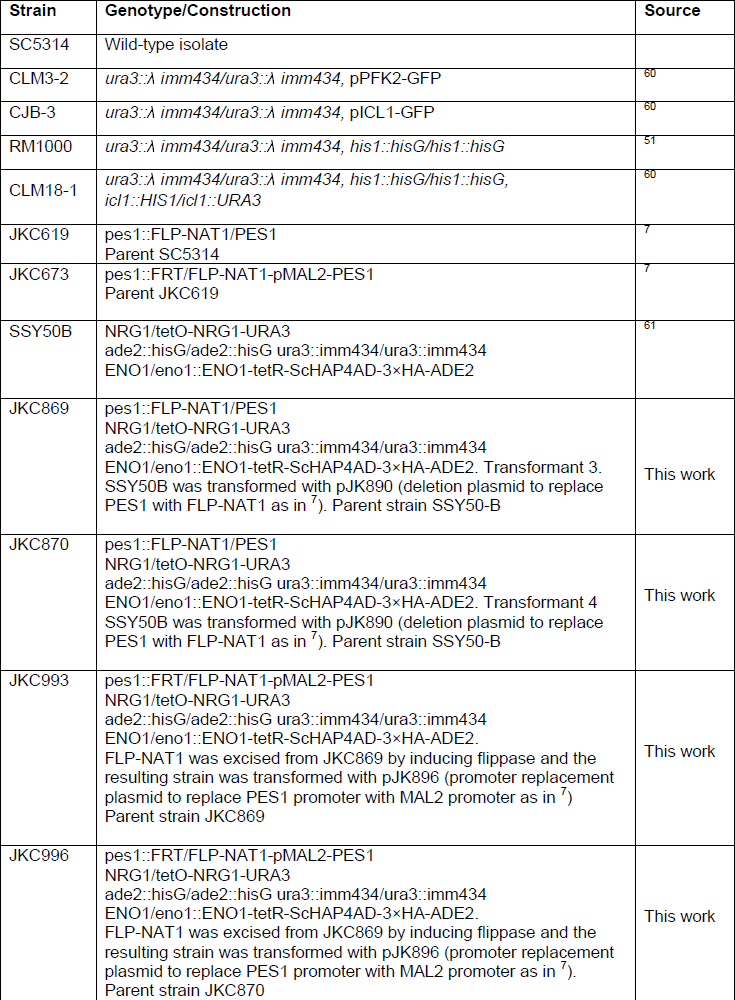
List of strains used in this study

#### Construction of the *TLOb2* KO strain

Strain 12822 (Haploid strain from Yue Wang (IMCB), Singapore) was transformed with the PCR product of pGEM-URA3 using primers^51^

5’TTTATGTTGTGCGAGACGTACTGTAATTTTATTGGTTGTGTGAGACCGGTTCTCGCCTTCCCAAATAGTGTGGAATTGTGAGCGGATA 3’ and 5’AACAAGAATAATTTTATCACACCTTTATTTCCCCCCACCATGCCAGAAAACCTCCAAACAAGATTACATAGTTTTCCCAGTCACGACGTT 3’. Transformants were selected for on SDC-ura and confirmed with primers 5’ AGACCTATAGTGAGAGAGCA 3’ / 5’ ACTTGTTGGACTTCGGAGTC 3’. This strain auto-diploidized based on flow cytometry and PCR so a second round of transformation to replace the second allele was performed. The heterozygous *TLOβ2/tloβ2* strain was transformed with a PCR product using primers 5’AACAAGAATAATTTTATCACACCTTTATTTCCCCCCACCATGCCAGAAAACCTCCAAACAAGATTACATAAGACCTATAGTGAGAGAGCA 3’ and 5’TTTATGTTGTGCGAGACGTACTGTAATTTTATTGGTTGTGTGAGACCGGTTCTCGCCTTCCCAAATAGGTAAAACGACGGCCAGTGAATTC 3’ from pGFP-NAT1^52^. Transformants were selected for on YPAD+Neurseothricin and confirmed with primer pair 5’ ACTTGTTGGACTTCGGAGTC 3’ and 5’ CAGATGCGAAGTTAAGTGCG 3’. PCR with *TLOb2* primers (5’ GATTACATAACTCACTCGACG 3’ and 5’ TACGTGGTTGGTGTTGTTGCCA 3’) confirmed that no wild-type copy remained.

### Cell collection and RNA extraction

Biofilms were developed in a simple flow biofilm model as described previously^12,13^. YNB+1% Glucose media was used as the medium for biofilm growth, with flow rate maintained at 1 ml/min. Biofilms were allowed to grow for 24 h. The biofilms were collected from the SE strip and flash frozen in liquid nitrogen followed by short-term storage at -80°C.

Prior to harvesting the biofilms, cells released from the biofilm in the flow-through media were collected at the completion of 24 h of biofilm formation. From our previous studies we have known that biofilms tend to disperse at the rate of ∼5x10^5^ – 1x10^6^ cells/ml in YNB medium^9^. To obtain enough cells for RNA extraction, we collected the flow-through several times from replicate biofilms, 3 ml at a time in tubes placed on ice. The cells were pelleted by cold ultracentrifugation, and the total time of collection of dispersed cells to freezing the pellet on dry ice and -80°C was 8 minutes. For some experiments, dispersed cells were enumerated by hemocytometer and by colony counts on solid medium (YPD-agar plates).

Planktonic cells growing for 24 h in flasks containing YNB+2% glucose were centrifuged and pellets stored at -80°C, for subsequent RNA extraction. RNA was extracted from all frozen pellets using a RiboPure yeast kit^22,53^.

#### Static biofilm formation

Biofilms were also grown under static conditions in microtiter plates as previously described^54^, with slight modifications. Briefly, one ml of *C. albicans* cells (1x10^6^ cells/ml) was introduced into wells of a 24-welled microtiter plate, incubated overnight in YNB broth plus or minus DOX, and in glucose or maltose as the carbon source. Next day, biofilms were washed twice and metabolic activity measured by using a semi-quantitative colorimetric 2,3-bis(2-methoxy-4-nitro-5-sulfophenyl)-2H-tetrazolium-5-carboxanilide (XTT) reduction assay as reported previously by our group ^54^. Biofilm aliquots were also visualized by light microscopy.

Biofilms were additionally developed on catheter material, as previously published^42^. Briefly, silicone elastomer (SE) pieces adhered with *C. albicans* were allowed to develop a biofilm in Spider medium contained in 6 well plates, at 37°C, for 48 h. The static biofilms were measured by XTT assay and dry weight, ^54^. For dry mass measurement of biofilms, medium was removed, biofilms disrupted, and the contents of each well were vacuum filtered over a pre-weighed 0.8 μm nitrocellulose filter (Millipore, AAWG02500). A control well with no cells added was also vacuum filtered. The filters were dried overnight, and weighed. The average total biomass for each biofilm was calculated from three independent samples after subtracting the mass of the filter with no cells. Statistical significance (p values) was calculated with a Student's one-tailed paired t test.

#### RNA sequencing and differential expression analysis

RNA was extracted from a total of six samples: two replicates each of planktonic, biofilm and biofilm dispersal cells. The RNA samples were sent to The Genome Sequencing Facility of Greehey Children's Cancer Research Institute at University of Texas Health Science Center at San Antonio, where the sequencing was executed, and bioinformatic analysis performed. Briefly, 5μg total RNA was used for mRNA isolation using Dynabeads OligodT (Invitrogen, Carlsbad, CA), then about 30-50ng isolated mRNA was used for mRNA-Seq library preparation by following the BIOO Scientific NEXTflex Directional RNA-Seq Kit (dUTP-based) sample preparation guide. The first step in the workflow involves purifying the poly-A containing mRNA molecules using Dynabeads poly-T oligo-attached magnetic beads. Following purification, the mRNA is fragmented into small pieces using divalent cautions under elevated temperature (95°C with 10 mins). The cleaved RNA fragments are copied into first strand cDNA using reverse transcriptase and random primers. This is followed by second strand cDNA synthesis using DNA Polymerase I and RNase H. These cDNA fragments then go through an end repair process, the addition of a single ‘A‘ base, and then ligation of the adapters. The products are then purified and enriched with PCR to create the final RNA-Seq library. Directionality is retained by adding dUTP during the second strand synthesis step and subsequent cleavage of the uridine containing strand using Uracil DNA Glycosylase. After RNA-Seq libraries were subjected to quantification process, pooled for cBot amplification and subsequent 100bp paired end sequencing run with Illumina HiSeq 2000 platform (San Diego, CA). After the sequencing run, demultiplexing with CASAVA was employed to generate the fastq file for each sample. All sequencing reads were filtered, trimmed and aligned with *C. albicans* reference genome using TopHat2^55^ default settings, and the Bam files from alignment were processed using HTSeq-count^56^ to obtain the counts per gene in all samples. A statistical analysis of differential gene expression was performed using DESeq package from Bioconductor^57^, and a gene was considered significantly altered if the false discovery rate for differential expression was ≤ 0.05.

The processes associated with differentially expressed genes were identified using Gene Ontology Slim Mapper^58^

#### Analysis of cellular DNA by fluorescence flow cytometry

Aliquots of dispersed cells collected from biofilms at 12 h, 24 h and 48 h were sonicated for 5 s and fixed overnight at 4°C in 95% ethanol. The cells were washed with 50 mM sodium citrate pH 7.0, suspended in the same buffer, and treated with 25 μl 10 mg ml^−1^ RNase A, and then with 25 μl of 20 mg ml^−1^ Proteinase K and incubated for 1 h at 50°C for both treatments. Finally, 1 ml of propidium iodide (PI) was added at a final concentration of 16 μg ml^−1^ and the samples were stored at 4°C. A total of 1 × 10 ^5^ PI stained cells were analyzed with a BD LSR II Flow Cytometer (Becton Dickinson, Franklin Lakes, NJ).

### Confocal Scanning Laser Microscopy

Biofilms were stained with 25 μg/ml Concanavalin A-Alexa Fluor 594 conjugate (C-11253; Molecular Probes, Eugene, Oregon, United States) for 1 hr in the dark at 37°C. Confocal scanning laser microscopy (CSLM) was performed with a Zeiss LSM 510 upright confocal microscope. Concanavalin A conjugate staining was observed using a HeNe1 laser with an excitation wavelength of 543 nm. For GFP visualization, an argon laser was used with 458-, 488-, and 514-nm excitation wavelengths. Images were overlaid and assembled into side and depth views using the Zeiss LSM Image Browser v4.2 software. In some studies, portions of biofilms were imaged only under bright-field, and/or these images were overlaid with the GFP 488 fluorescence of the same cell population.

### Epistasis study under planktonic conditions

Cells were grown on Yeast Extract 2% Maltose (YPM) plates containing 5 ng/ml doxycycline (DOX) for 48 h at 30°C. After washing twice with 0.9% NaCl, five fold cell dilutions beginning at OD_600_ of 0.5 were spotted with a caliberated replicator onto YPM, Yeast Extract + 2% Dextrose (YPD), Spider without mannitol with 2% Maltose (SpiM) or with 2% Dextrose (SpiD). Plates contained 5 µg/ml DOX or vehicle. Plates were incubated for 3 days at 30 °C (YP) or 37°C (Spi).

### *In vivo* jugular vein catheter model

A *C. albicans* mouse catheter biofilm model was used for *in vivo* experiments as previously described^59^ with slight modifications. Following receipt of the jugular vein catheterized mice from Charles River labs (Wilmington, MA), the catheters were instilled with 25 µl of *C. albicans* inoculum of 5x10^6^ cells/ml (entire catheter volume). Two strains were used – the wildtype and *pes1/PES1-tet*, and were first germinated for 90 minutes in the absence of DOX in YNB medium. The mice were divided into three sets of 5 mice each, a) mice in which catheters were instilled with the wildtype strain in YNB, b) mice catheters inoculated with *pes1/PES1-tet* in YNB, and c) catheters instilled with pes1*/PES1-tet* in YNB containing 25 µM doxycycline. The cells were allowed to dwell in the catheters for 3 days, after which the mice were sacrificed and catheters aseptically removed. The catheters were cut laterally and imaged under a phase contrast microscope to visualize the morphology of the cells growing inside the catheters of the individual groups. Additionally, the kidneys of the sacrificed mice were harvested, homogenized and plated on YPD to enumerate the extent of fungal burden in the organ. Differences in the organ burden between the three groups were tabulated as Log cfu and a two tailed t-test with a p value <0.05 was considered significant.

---

**Supplemental data 1 (S1). Comprehensive differentially expressed data of >1.5 fold significantly (p<0.5) regulated genes.** In this excel sheet, Sheet1: Genes significantly upregulated in biofilm versus planktonic cells; Sheet 2: Genes significantly downregulated in biofilm versus planktonic cells; Sheet 3: Genes common between our study versus other transcriptone studies comparing biofilm to planktonic expression data. Sheet 4: Genes significantly upregulated in dispersed versus planktonic cells; Sheet 5: Genes significantly downregulated in dispersed versus planktonic cells; Sheet 6: Genes showing similar patterns of upregulation in both biofilm and planktonic, compared to planktonic cells; Sheet 7: Genes showing similar patterns of downregulation in both biofilm and planktonic, compared to planktonic cells.

**Supplemental data 2 (S2). Dataset of genes expressed in dispersed cell population.** In this excel sheet, Sheet 1: Differential expression of Central carbon metabolism genes in dispersed and planktonic cells, both versus biofilm cells. Here the dispersed cells show planktonic yeast-like expression pattern. Sheet 2: Set of genes that are upregulated and downregulated “only” on dispersed cells. This was extracted after pair-wise comparisons of all the sample conditions to eachother, and picking largely that were up or down exclusively in dispersed cells.

**Supplemental data 3 (S3). Slide1: Figure is related to figure 2 in the manuscript**. This shows that the topmost layers of the flow biofilm expressis green fluorescent Pfk2. Slide 2: Figure is related to figure 2 in the manuscript. This shows that the cells attached to the silicone substrate (innermost portion of the flow biofilm) express green fluorescent Icl1.

